# PGxO and PGxLOD: a reconciliation of pharmacogenomic knowledge of various provenances, enabling further comparison

**DOI:** 10.1101/390971

**Authors:** Pierre Monnin, Jöel Legrand, Graziella Husson, Patrice Ringot, Andon Tchechmedjiev, Clément Jonquet, Amedeo Napoli, Adrien Coulet

## Abstract

**Background:** Pharmacogenomics (PGx) studies how genomic variations impact variations in drug response phenotypes. Knowledge in pharmacogenomics is typically composed of units that have the form of ternary relationships *gene variant – drug – adverse event*. Such a relationship states that an adverse event may occur for patients having the specified gene variant and being exposed to the specified drug. State-of-the-art knowledge in PGx is mainly available in reference databases such as PharmGKB and reported in scientific biomedical literature. But, PGx knowledge can also be discovered from clinical data, such as Electronic Health Records (EHRs), and in this case, may either correspond to new knowledge or confirm state-of-the-art knowledge that lacks “clinical counterpart” or validation. For this reason, there is a need for automatic comparison of knowledge units from distinct sources.

**Results:** In this article, we propose an approach, based on Semantic Web technologies, to represent and compare PGx knowledge units. To this end, we developed PGxO, a simple ontology that represents PGx knowledge units and their components. Combined with PROV-O, an ontology developed by the W3C to represent provenance information, PGxO enables encoding and associating provenance information to PGx relationships. Additionally, we introduce a set of rules to *reconcile* PGx knowledge, i.e. to identify when two relationships, potentially expressed using different vocabularies and levels of granularity, refer to the same, or to different knowledge units. We evaluated our ontology and rules by populating PGxO with knowledge units extracted from PharmGKB (2,701), the literature (65,720) and from discoveries reported in EHR analysis studies (only 10, manually extracted); and by testing their similarity. We called PGxLOD (*PGx Linked Open Data*) the resulting knowledge base that represents and reconciles knowledge units of those various origins.

**Conclusions:** The proposed ontology and reconciliation rules constitute a first step toward a more complete framework for knowledge comparison in PGx. In this direction, the experimental instantiation of PGxO, named PGxLOD, illustrates the ability and difficulties of reconciling various existing knowledge sources.

## Background

In this article, we present a simple ontology named PGxO, developed to reconcile and trace knowledge in pharmacogenomics (PGx). We instantiated this ontology with knowledge of various origins to both illustrate the relevance of the ontology and constitute a Linked Open Data (LOD) data set for PGx [5].

PGx itself studies how genomics impact individual variations in drug response phenotypes [63]. Knowledge in pharmacogenomics is of particular interest for the implementation of personalized medicine, i.e. a medicine tailoring treatments (chosen drugs and dosages) to every patient, in order to reduce the risk of adverse events. Indeed, best known examples of PGx knowledge already led to the development of clinical guidelines and practices [9] that recommend considering individual genotype when prescribing some particular drugs such as abacavir (an anti-retroviral) or fluorouracile (an anti-neoplastic) [2, 35].

Units of PGx knowledge typically have the form of a ternary relationship *gene variant – drug – adverse event* stating that a patient having the gene variant and being treated with the drug will have a higher risk of developing the mentioned adverse event. For example, one relationship is *G6PD:202A – chloroquine – anemia*, stating that patients with the 202A version of the G6PD gene and treated with chloroquine (an anti-malarial drug) are likely to experience anemia (an abnormally low level of red blood cells in blood).

An increasing volume of state-of-the-art knowledge in PGx can be found in reference databases, such as PharmGKB [61], or in the biomedical literature [21]. But, a large part of this knowledge is suffering from a lack of validation, or “clinical counterpart” [26], and is not yet ready to be translated into clinical guidelines and practices. For example, about 91% (as of July 2018) of the relationships listed in PharmGKB are qualified with a level of evidence 3 or 4, corresponding, at best, to an unreplicated study or to multiple studies that show a lack of evidence for the relationship [61]. On the other hand, PGx knowledge can also be discovered from clinical data, such as Electronic Health Records (EHRs), particularly when those are linked to DNA biobanks [4, 14, 49]. In this case, discovered knowledge can either be new or can interestingly temper, or confirm, knowledge elsewhere stated, but that may lack validation.

However, comparing PGx knowledge coming from distinct sources is challenging because of the heterogeneity of these sources. Indeed, such sources may use different vocabularies, different levels of granularity or even different languages to represent knowledge units. Therefore, there is a strong need for developing a common schema that would enable comparing knowledge extracted or discovered from various sources. Several ontologies have already been developed for pharmacogenomics, but with different purposes, making them unadapted to the present need. In particular, SO-Pharm (Suggested Ontology for Pharmacogenomics) and PO (Pharmacogenomic Ontology) have been developed for knowledge discovery purposes rather than data integration or knowledge reconciliation [12, 17]. The PHARE ontology (for PHArmacogenomic RElationships) has been built for normalizing *gene — drug* and *gene — disease* relationships extracted from text and is not suitable for representing ternary PGx relationships [11]. More recently, Samwald *et al.* introduced the Pharmacogenomic Clinical Decision Support (or Genomic CDS) ontology, whose main goal is to propose consistent information about pharmacogenomic patient testing to the point of care, to guide physician decisions in clinical practice [54]. We have built PGxO by learning and adapting from these previous experiences. For consistency reasons and good practices, we mapped PGxO to concepts of these four pre-existing ontologies.

In this work, we propose to leverage Semantic Web and Linked Open Data (LOD) [5] technologies as a first step toward building a framework to represent and compare PGx relationships from various sources. We import knowledge of three origins to instantiate our “pivot” ontology, both illustrating the role of the ontology, and building a community resource for PGx research.

In the preliminary stage of this work [37], we proposed: *(i)* a first version of the PGxO ontology able to represent simple pharmacogenomic relationships and their potentially multiple provenances and *(ii)* a set of rules to reconcile PGx knowledge extracted from or discovered in various sources, i.e. to identify when two relationships refer to the same, or to different knowledge units. In this paper, we extend PGxO to improve its ability to represent PGx relationships extracted from the literature and by adding the notion of *proxy*. We experiment our approach by instantiating PGxO with knowledge of various provenances: PharmGKB, the biomedical literature, and results of studies that analyzed EHR data and linked DNA biobanks [4].

This paper is organized as follows. The *Methods* section introduces the methods used for the construction of PGxO, for encoding provenance information and for our rule-based approach to reconcile PGx knowledge. Details are also given about algorithms and techniques used to instantiate PGxO from the aforementioned sources. The *Results* section presents our ontology, PGxO, the reconciliation rules and results of the instantiation and reconciliation processes. Finally, we conclude this work by discussing the abilities of PGxO for representing and reconciling PGx knowledge and the next challenges to tackle.

## Methods

### Ontology construction

PGxO was manually and collaboratively developed by 3 persons (PM, CJ and AC) in 7 iterations (as of July 2018). We followed classical ontology construction methods and life cycle [16, 41], including the steps of specification, conception, diffusion and evaluation of the ontology.

#### Specification

Our aim is to represent and reconcile what we defined as PGx knowledge units. These are ternary relationships between one (or more) *genetic factor(s)*, one (or more) *drug treatment(s)* and one (or more) *phenotype(s)*. Such phenotypes can be the expected outcomes of the drug treatments or some adverse effects. In order to keep our ontology simple and leverage existing works representing PGx components, we restrain the *scope* of PGxO only to representing PGx knowledge units and not all facets of pharmacogenomics. The *objective* of PGxO is twofold: reconciling and tracing these PGx knowledge units. To enable this reconciliation, we need to encode metadata and provenance information about a PGx relationship.

#### Conception and Diffusion

Because PGxO is of small size, the conception step was performed simultaneously with conceptualization, formalization and implementation steps. The ontology has been implemented in OWL using the Protégé ontology editor [39]. PGxO is conceived around the central class of PharmacogenomicRelationship, which enables associating two or three of the following key components of PGx: `Drug, GeneticFactor` and `Phenotype`. The expressive Description Logic (DL) associated with PGxO is 𝒜 ℒ 𝒞 𝓘 (*D*) [3]. Successive versions of PGxO have been published online and shared with collaborators through both the NCBO BioPortal [1, 42] and GitHub [47]. We have followed [36] guidelines to report on the Minimum Information for the Reporting of an Ontology (MIRO) associated with PGxO and made this available at [47].

#### Evaluation

To evaluate our ontology, we used *competency questions* as proposed by Gangemi [20]. The questions we defined are the following:

1. Does PGxO enable to represent a PGx knowledge unit from the PGx state of the art (i.e. from a reference database or extracted from the biomedical literature), along with its provenance?
2. Does PGxO enable to represent a PGx knowledge unit discovered from clinical data, along with its provenance?
3. Does PGxO, coupled with its reconciliation rules, enable to decide if two knowledge units, with distinct provenances, may refer to the same thing?

We answered these questions twice, once early and once late in the iterations of the development of PGxO. For the former iteration, we manually instantiated PGxO with examples of knowledge units, associated with their provenances, from *(i)* PharmGKB, *(ii)* the literature (extracted by Semantic Medline [52] or FACTA+ [57]) and *(iii)* hand designed facts corresponding to what we thought may be discovered in EHRs. For the latter iteration, we answered these questions by instantiating PGxO with knowledge units extracted programmatically from PharmGKB and the biomedical literature, and manually from results reported by studies analyzing EHR data and linked biobanks. Details on the methods used to populate PGxO from these various sources are provided in following subsections.

#### Mappings

For consistency reasons and good practices, we manually mapped concepts of PGxO to the four afore-mentioned ontologies related to pharmacogenomics: SO-Pharm, PO, PHARE and Genomic CDS. These mappings are available in [46]. Because the NCBO BioPortal generates lexical-based mappings between the ontologies it hosts, it provides an initial set of mappings from PGxO to many standard ontologies. In particular, we manually completed PGxO BioPortal mappings to three standard and broad spectrum ontologies: MeSH, NCIt and SNOMED CT. These mappings are available in [45]. The two resulting sets of mappings are provided as independent OWL files to allow a flexible loading of the ontology with, or without mappings.

### Provenance encoding

Data provenance (sometimes called lineage) refers to metadata that state where data came from, how they were derived, manipulated, and combined, and how they may have been updated [7]. With PGxO, we do not only want to represent units of knowledge of different origins, but also to trace their origins. Therefore, we need an encoding of the provenance of knowledge units. For this purpose, we leverage an existing ontology for provenance, named PROV-O [33], which is a W3C Recommendation since 2013. In addition, for some particular provenance metadata, PGxO reuses object properties of the high-level ontology DUL (Dolce+DnS Ultralight) [19].

PROV-O is built around three main concepts: `Entity`, `Activity` and `Agent`. Entities represent things that can be generated, modified, etc. by activities. Activities are realized by agents that can be either human or software agents. Entities can also be directly attributed to agents.

In terms of PROV-O concepts, authorities which publish sources from which we extract knowledge units are considered to be agents (e.g. the PharmGKB team, the National Library of Medicine (NLM) in charge of PubMed, an hospital in charge of a repository of EHRs). Data sources (e.g. a version of PharmGKB, of PubMed, a repository of EHRs) are attributed to agents, and then may be used to derive data. These data, in turn, are used during the execution of an activity (e.g. a mining algorithm). Such execution generates entities that in our case are PGx knowledge units. Quantitative and qualitative metadata may be associated to an activity and to the entities it generates. For instance, one can specify the version of an algorithm, the date of its execution, the quality of the generated entities (such as their levels of confidence, their p-values or odds ratios). Thereby, a further comparison of two knowledge units having different or identical provenances may take into account these various elements.

### Reconciliation rules

Besides representing and encoding heterogeneous PGx relationships within a single knowledge base, a comparison mechanism is needed to identify when two relationships refer to the same knowledge unit or not. However, the description of PGx relationships is highly heterogeneous depending on the sources we consider, leading to many relationships being similar to some extent, but not exactly identical. For example, one source may document a relationship between a gene variant *gv*_1_, a drug *d*_1_ that causes a drug response phenotype *p*_1_, whereas a second source may only document the relationship between *gv*_1_ and *d*_1_, omitting any drug response phenotype. Alternatively, a third source may document the same relationship at a broader level, for instance by mentioning only the involved gene *g*_1_, instead of stating the causative variant *gv*_1_ (that is part of *g*_1_).

To take into account this variability, we defined a set of rules enabling basic comparison mechanisms. The rules focus on identifying identical relationships, broader/narrower ones and related ones (to some extent). They compare two PGx relationships using their associated components (i.e. drugs, genetic factors, phenotypes). To represent and implement the defined rules, we investigated semantic web rule languages, such as SWRL and DL-Safe rules [23,24,30]. Unfortunately, we found them unadapted to our case, for expressiveness reasons. Indeed, those formalisms do not allow to check equalities or inclusions of sets of individuals, which instantiate defined DL expressions (see Additional file 1 for examples). Therefore, we represent the rules with our own formalism. On the left side of a rule, equalities or inclusions between sets of components of the two compared PGx relationships are tested and return a boolean result. Tests of equalities or inclusions can be combined using conjunctions (AND) and / or disjunctions (OR). On the right side, a link between the PGx relationships is to be added to the populated ontology if and only if the left side of the rule is true. This link can specify the two PGx relationships as identical, related or one being broader than the other. As the rules are not formalized using a particular semantic web rule language, they are kept separated from the definitions of PGxO. Therefore, we implemented them in an independent Python program interacting with the populated ontology using the SPARQL query language. This program is executed once, derives novel facts from the rules and adds them to the knowledge base.

We take advantage of Semantic Web technologies, by using associated reasoning mechanisms for the comparison of PGx relationships. In particular, we use the semantics associated with `owl:sameAs` links that state that two URIs are actually referring to the same entity. The interpretation of the `rdfs:subClassOf relation` and its transitivity is also used when comparing a PGx relationship that may be more specific/general than another one.

### Ontology instantiation

We instantiated the ontology with PGx knowledge units from various sources, first, to answer the *competency questions* defined for PGxO and, second, to build a data set called PGxLOD (*PGx Linked Open Data*) that we think may constitute a valuable community resource for PGx research.

#### With preexisting LOD

We initiated PGxLOD from a preexisting set of Linked Open Data made of interconnected genes, drugs, and diseases according to 6 standard databases [13]. Such LOD data set follows the Semantic Web standards, particularly by using the Resource Description Framework (RDF) language and Uniform Resource Identifiers (URI) [5], which makes it adequate to connect with Semantic Web ontologies.

This preexisting data set is an aggregation of data from ClinVar [32], DisGeNET [43], DrugBank [62], SIDER [31] and MediSpan (a proprietary database). Accordingly, it includes and relates data about drugs, diseases and phenotypes, but no PGx relationships. Nevertheless, this data set groups together data related to entities involved in PGx relationships, and mappings between entities that may be present in different data sources, e.g. a drug referenced both in DrugBank and SIDER. These mappings are of particular interest for our purpose of comparing PGx relationships, since those may be composed of entities referenced in these various sources.

The initial instantiation of PGxO with preexisting LOD is straightforward since entities representing genes, drugs and diseases are used to instantiate the corresponding PGxO concepts, using the RDF predicate `rdf:type`. In the following, we name PGxLOD v1 the result of this instantiation process. This constitutes the initial version of PGxLOD, with no PGx relationships, to distinguish from PGxLOD v2, a version enriched with PGx relationships of various provenances.

#### With PharmGKB data

Second, PGxO was instantiated with data from PharmGKB [61], a reference database for pharmacogenomics. PharmGKB’s *clinical annotations* describe PGx relationships between genes (potentially their variants), drugs, and phenotypes. They are produced by PharmGKB curators after a review of the biomedical literature and of recommendations of health agencies such as the US Food and Drug Administration. In addition, PharmGKB contains cross-references, i.e. identifiers of genes, variants, drugs and phenotypes within other databases (such as NCBI Gene for genes) or ontologies (such as the Anatomical Therapeutic Chemical Classification System for drugs).

Part of PharmGKB data are already available in the form of LOD as a part of the Bio2RDF project [8]. Nonetheless, this version is outdated and provides only a small portion of PharmGKB. Therefore, inspired by this precursor work and following the guidelines of the Bio2RDF project, we developed new scripts producing a more complete RDF version of PharmGKB. These scripts transform the latest downloadable text files of PharmGKB, first, into a SQL database (with a script named *pharmgkb2sql*) and then into RDF triples (with a script named *pharmgkbsql2triples*).

Drug response phenotypes provided in clinical annotations by PharmGKB are non easy to translate as they are reported within plain-text sentences. Because PharmGKB also provides a broad type of drug responses (Efficacy, Toxicity/ADR, Metabolism/PK, Dosage, Other) in a structured manner, we decided, for simplicity, to use those directly instead of further text mining analysis on textual descriptions, and then considered only Efficacy and Toxicity/ADR, because they encompass the drug response phenotypes we want to capture. Accordingly, PGx relationships in PGxLOD are associated with these two high level types of drug responses.

Components of PGx relationships (i.e. drugs, genes and variants) are represented with URIs using both PharmGKB identifiers and Bio2RDF naming conventions. In addition, PharmGKB cross-references to external databases and ontologies are used to map PharmGKB URIs either to URIs already defined in our LOD, or to new ones. The type of relation used is `bio2rdf:x-ref` for every cross reference; doubled with either a `owl:sameAs` relation when the cross-reference points to an identical entity in an external database, or with a `rdf:type` relation when it points to an ontology class.

Among the metadata associated with PharmGKB clinical annotations, we particularly keep the *level of evidence*. Levels of evidence have been defined by PharmGKB [61] as a six-level scale (1A, 1B, 2A, 2B, 3, 4), where higher levels (1A, 1B, 2A, 2B) indicate that a relationship has been significantly studied or is medically implemented; and lower levels (3, 4) indicate that the PGx relationship has only been reported in a single study or lacks clear evidence. Levels of evidence are of particular importance as they may help us identify PGx knowledge with irregular levels of validation in various sources.

#### With the biomedical literature

Third, we instantiated PGxO with elements extracted automatically from the biomedical literature. Here we used a supervised machine learning prototype for relation extraction from text, trained on a manually annotated corpus. Please note that in this work, we are not aiming at achieving the best performance, but rather aiming at showing that we can extract PGx relationships from text, along with their provenance metadata, and compare these to others, extracted from distinct sources. This illustrates that PGxO enables structuring, documenting (the name of the algorithm used, its performance, etc.), then comparing a relationship extracted from text.

We assembled a set of 657,538 sentences from 86,520 PubMed abstracts related to pharmacogenomics. Removing malformed sentences, based on tokenization errors, and sentences that do not contain at least one drug and one genetic factor, based on named entities recognized by PubTator [59], we obtained a corpus of 176,704 sentences. Out of those, we randomly selected 307 sentences and had each sentence manually annotated with the BRAT software [56], by 3 distinct annotators from a group of 11 pharmacists, biologists and bioinformaticians. The annotation task is precisely specified in annotation guidelines, publicly available [44]. Annotations are limited here to four broad entity types, mainly involved in PGx relationships: Genes, Genomic Variations, Drugs and Phenotypes and to two broad relation types, “isAssociatedWith” and “isEquivalentTo”, between these entities. The latter is used only to relate the numerous acronyms to their extended form. An example of an annotated sentence is shown in Figure 1, and the main characteristics of the corpus are summarized in Table 1.

**Figure 1.**
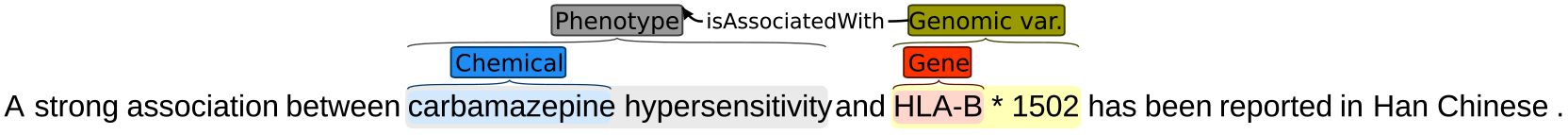
Example of a sentence (PMID=18370849), manually annotated with four entities and one relation.

**Table 1.** Statistics of named entities and relations manually annotated in our 307-sentence corpus. Entities annotated in multiple sentences are counted multiple times. Second-layer entities refer to entities which offset includes the annotation of a first-layer entity

| Named entities |  |  | Relations |  |
| --- | --- | --- | --- | --- |
| Type | First-layer | Second-layer | Type |  |
| Gene | 452 | 20 | isAssociatedWith | 582 |
| GenomicVariation | 74 | 166 | isEquivalentTo | 77 |
| Drug | 459 | 36 |  |  |
| Phenotype | 262 | 251 |  |  |
| Total |  | 1720 | Total | 659 |

Our approach is classically composed of two steps: a Named Entity Recognition (NER), followed by a relation extraction. Two supervised machine learning models were trained for the first step, and a third one for the second step. The NER is performed using a Convolutional Neural Network (CNN) model described in [10], trained on the 307 annotated sentences. We first annotate shallow entities (Figure 1) using an instance of this model with *PubTator* annotations as an input. We name these entities *first–layer* entities. Then, a second instance of the same model is trained to annotate *second layer* entities, i.e. entities with an offset that includes a *first layer* entity, using the *first layer* entities as input. In Figure 1, *first layer* entities would be *carbamazepine* and *HLA B*, and *second layer* entities are ***carbamazepine*** *hypersensitivity* and ***HLA-B****\*1502*. Finally, we trained a model similar to the one described in [48] to extract relationships between identified entities.

Reasonable meta-parameters were selected according to previous experiments. The size of the word embeddings was set to 100. The size of the *PubTator* and *first layer* embeddings was set to 20. The kernel size of the convolution was set to 100. Word embeddings were pre-trained on ~3.4 million PubMed abstracts (corresponding to all those published between Jan. 1, 2014 and Dec. 31, 2016) using the method described in [34]. Performances of the two steps, evaluated on a 10-fold cross validation, are respectively reported in Table 2 and 3. In these two tables, the f1-score is defined as the harmonic mean of the precision and recall, i.e. 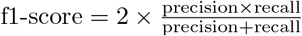.

**Table 2.** Named Entity Recognition (NER) performance in terms of precision (P), recall (R) and f1-score (F1). Results of second-layer entities take into account the prediction error of the first-layer entities. Std stands for F1 standard deviation.

|  | P | R | F1 | std |
| --- | --- | --- | --- | --- |
| Drug | 0.92 | 0.87 | 0.89 | 0.03 |
| Gene | 0.97 | 0.91 | 0.94 | 0.03 |
| Phenotype | 0.84 | 0.66 | 0.74 | 0.09 |
| Genomic variation | 0.81 | 0.69 | 0.74 | 0.08 |
| All entities | 0.86 | 0.80 | 0.83 | 0.05 |

**Table 3.**
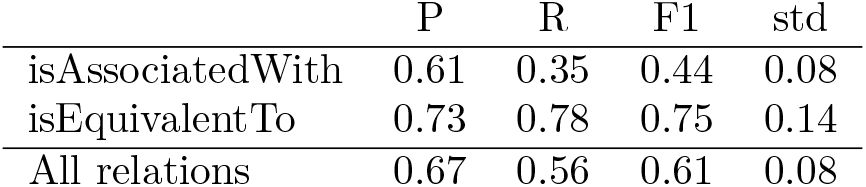
Relation extraction performance in terms of precision (P), recall (R) and f1-score (F1). Std stands for F1 standard deviation. Results take into account the prediction error for the entities.

|  | P | R | F1 | std |
| --- | --- | --- | --- | --- |
| isAssociatedWith | 0.61 | 0.35 | 0.44 | 0.08 |
| isEquivalentTo | 0.73 | 0.78 | 0.75 | 0.14 |
| All relations | 0.67 | 0.56 | 0.61 | 0.08 |

Trained models are applied to a test set of 176,704 sentences of PubMed abstracts, to extract automatically relations from text. After filtering out relationships that relate two GenomicVariations, two Phenotypes, or two Drugs, both the manually annotated relations and the automatically extracted ones are transformed into RDF.

Types of entities are manually aligned to the corresponding classes of PGxO. Each annotated and extracted entity is associated with an URI that is constructed, depending on its type, either from an identifier of a reference database (such as NCBI Gene for genes) or from an identifier of an ontology (such as ATC for drugs). Distinct reference databases or ontologies may be used for each type of entities depending on their availability. Accordingly, we defined an arbitrary order of choice for searching for references, presented in Table 4. This choice was motivated in part by the output of *PubTator*. For each type, the procedure is the following: given an entity, the first reference database or ontology is searched for the entry using string matching; if no entry matches, the next reference database or ontology is searched. Lastly, if no entry is found, we create a local URI within the PGxLOD namespace. Considering an extracted entity, when an entry is found in a reference database, its identifier is used to construct the corresponding URI. When an entry is found in an ontology (i.e. a class of an ontology), the extracted entity is given a local URI, and instantiates the ontology class.

**Table 4.** Reference databases and ontologies used to normalize the entities extracted from text. PGxLOD means that a local URI is created.

| Order | Drug | Gene | GenomicVariation | Phenotype |
| --- | --- | --- | --- | --- |
| 1<sup>st</sup> | MeSH | NCBI Gene | dbSNP | MeSH |
| 2<sup>nd</sup> | ChEBI | PGxLOD | PGxLOD | MEDDRA |
| 3<sup>rd</sup> | ATC |  |  | PGxLOD |
| 4<sup>th</sup> | PGxLOD |  |  |  |

#### With Electronic Health Records and linked biobanks studies

Fourth, we instantiated PGxO with PGx knowledge manually extracted from the reading of ten studies on patient Electronic Health Records (EHRs) and linked biobanks [4, 14, 18, 28, 29, 38, 40, 50, 58, 60]. For instance, Kawai *et al.* [29] report a statistical association (OR=2.05, 95%) between the haplotype CYP2C9*3, and severe bleeding, in patients treated with warfarin. Their study was performed on a biobank named BioVU, linked to patient EHRs of the Vanderbuilt University Hospital [53]. The ten studies were selected from mentions in CPIC (Clinical Pharmacogenetics Implementation Consortium) guidelines [9] and in the literature review of Denny *et al.* [15].

Entities involved in PGx relationships were manually associated with URIs already defined in PGxO and PGxLOD. The aim here is to assess the adequacy of PGxO to represent results of such studies. Indeed, it is one of our use cases to consider PGx researchers who want to compare the results they obtained on their local biobanks+EHRs, to results elsewhere reported.

## Results

### PGxO

Illustrated in Figure 2, PGxO is composed of eleven concepts (`owl:Class`), organized around the central concept `PharmacogenomicRelationship`, which represents an atomic unit of PGx knowledge.

**Figure 2.**
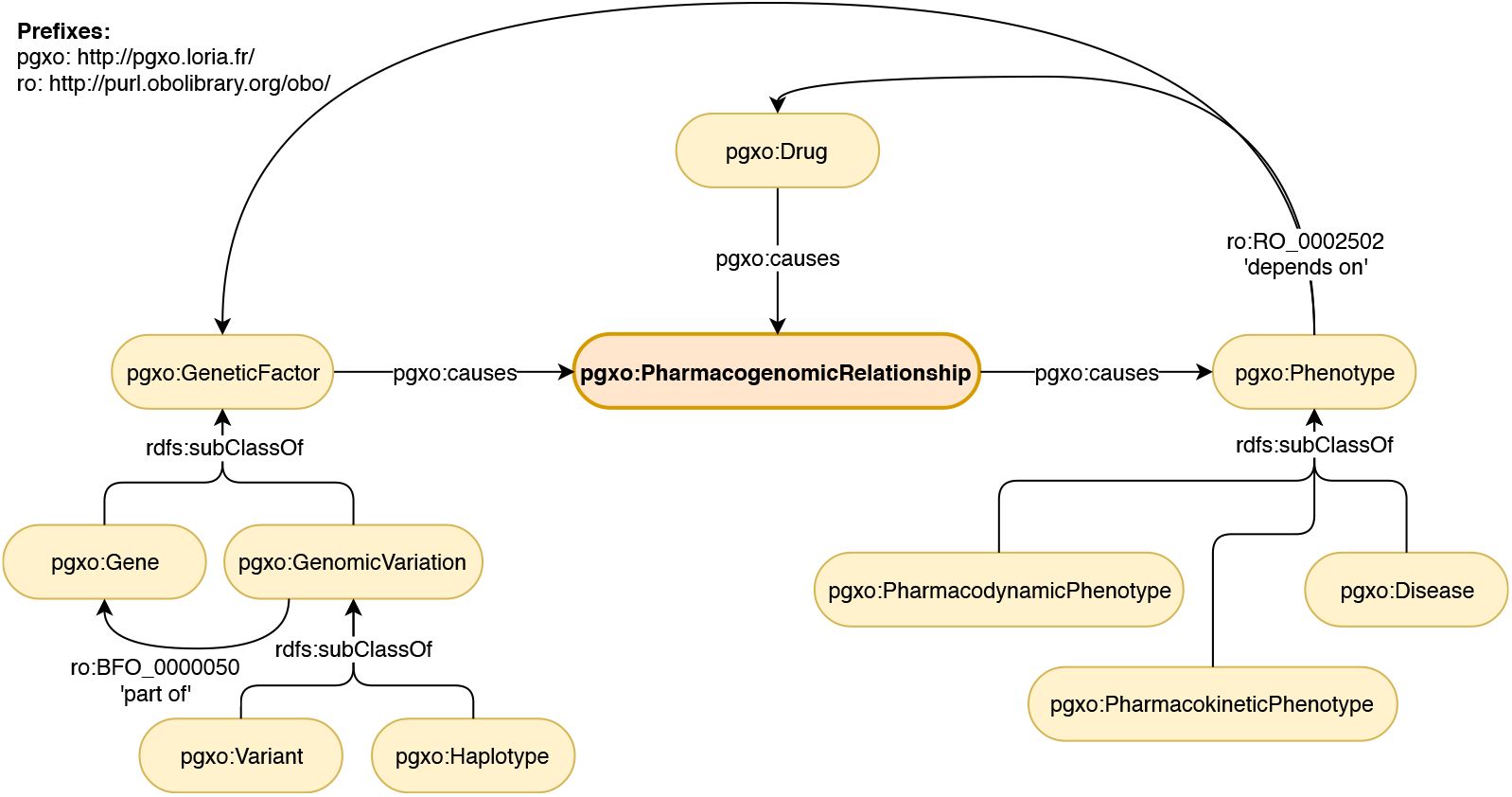
Main concepts and relations of PGxO. The central concept of the ontology is PharmacogenomicRelationship.

Instances of the concepts may be related by various types of relations (`i.e. owl:ObjectProperty`). Relation types causes and isCausedBy are used to connect components of a PGx relationship and are defined as inverses: causes ≡ isCausedBy^*−*^. The relation type `partOf (or ro:BFO 0000050)`, from the Relation Ontology (RO) [55], is used to express that a genomic variation is located within the sequence of a specific gene. The relation type `dependsOn (ro:RO 0002502)`, also from RO, is used to express complex phenotypes that involve other entities, e.g. gene expression such as *the expression of VKORC1* or drug response phenotypes such as *carbamazepine hypersensitivity*.

The concept `PharmacogenomicRelationship` is described by a Description Logics [3] axiom that may be decomposed into a union of three other axioms. Indeed, we consider that a PGx relationship involves at least two or three of the following components: drugs, genetic factors and phenotypes (i.e. drug response phenotypes). For more flexibility, we allow drug components to be either a drug, or something that depends on a drug (Axiom 1). Similarly, we allow genetic factor components to be either a genetic factor, or something that depends on a genetic factor (Axiom 2). This flexibility allows to represent relationships that involve, for instance, the expression of a gene (something that depends on a genetic factor, e.g. *the expression of VKORC1*) instead of a gene, or a drug resistance or sensitivity (something that depends on a drug, e.g. *carbamazepine hypersensitivity*) instead of a drug.

**Axiom 1.** DComponent *≡* Drug ⨆ ∃ dependsOn.Drug

**Axiom 2.** GFComponent *≡* GeneticFactor ⨆ ∃ dependsOn.GeneticFactor

Another flexibility resides in allowing a PGx relationship to have only two of these three components. Indeed, one component may be missing for multiple reasons: the relationship may still be under study and some of its components unknown; we can also expect errors during automatic extraction leading to missing components. Therefore, Axioms 3, 4 and 5 represent the three possibilities of having two components among the three considered. Axiom 6 is the final axiom describing the concept PharmacogenomicRelationship. A PGx relationship involving the three types of components trivially validates the axioms.

**Axiom 3.** PR_1_ ≡ ∃ causes.Phenotype ⨅ ∃ isCausedBy.DComponent

**Axiom 4.** PR_2_ *≡* ∃ causes.Phenotype ⨅ ∃ isCausedBy.GFComponent

**Axiom 5.** PR_3_ *≡* ∃ isCausedBy.DComponent ⨅ ∃ isCausedBy.GFComponent

**Axiom 6.** PharmacogenomicRelationship ⊑ PR_1_ ⨆ PR_2_ ⨆ PR_3_

Axiom 6 is not a concept definition (using ≡) because we consider that presenting two (or three) components is a required condition, but not a sufficient one to be a `PharmacogenomicRelationship`. Examples of encoding of PGx relationships and their provenance are detailed in the next subsection.

### PGxLOD: an instantiation of PGxO with knowledge units of various provenances

PGxLOD is the knowledge base that results from the instantiation processes of PGxO. Because a large portion of the data of PGxLOD has no license restriction, we provide an open access to this portion (only) at https://pgxlod.loria.fr. Full access to PGxLOD that includes the small set of proprietary data is granted upon request. The population processes of PGxLOD are detailed in the next paragraphs and their results are summarized in Table 5.

**Table 5.**
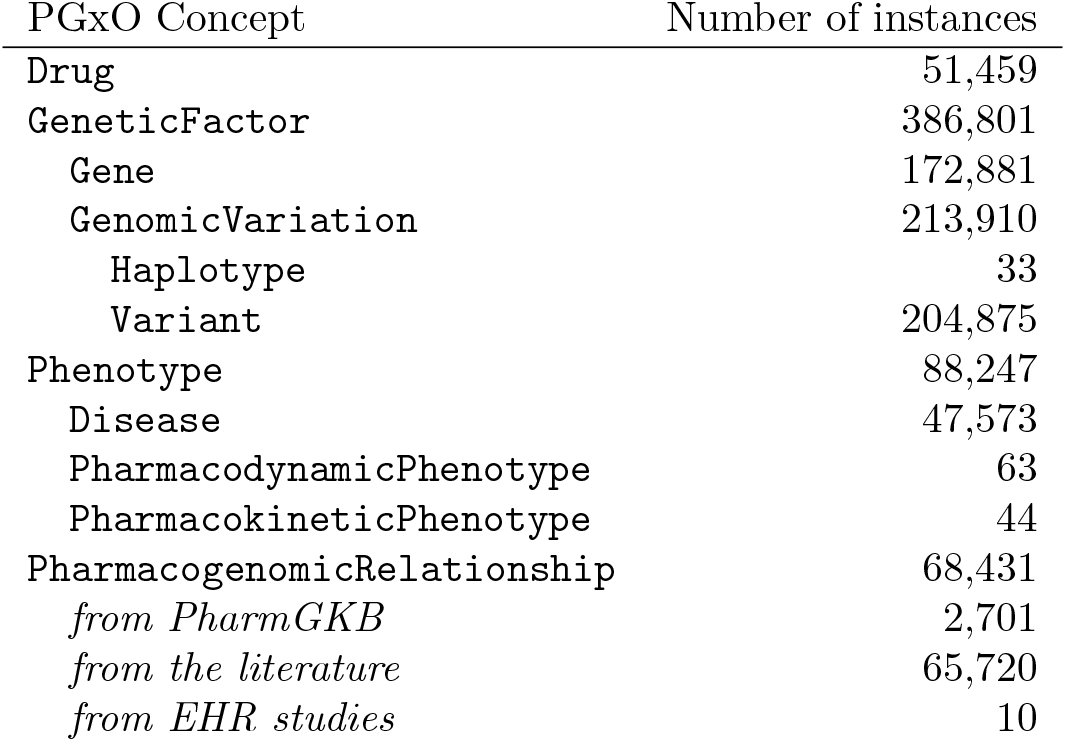
Main statistics of PGxLOD v2

#### With preexisting LOD

Table 6 summarizes results of the instantiation of PGxO with our preexisting LOD. At this stage PGxLOD does not contain any PGx relationship, but provides entities appearing as components of PGx relationships, as well as mappings between these entities.

**Table 6.** Statistics of the instantiation of PGxO with data from PGxLOD v1.

| Source | Genes | Variants | Drugs | Diseases | Phenotypes |
| --- | --- | --- | --- | --- | --- |
| ClinVar | 21,487 | 103,219 | 0 | 0 | 6,837 |
| DisGeNET | 85,893 | 49,279 | 0 | 38,727 | 6,092 |
| DrugBank | 4,300 | 0 | 7,740 | 0 | 0 |
| MediSpan | 0 | 0 | 5,820 | 2,481 | 0 |
| SIDER | 0 | 0 | 25,479 | 6,291 | 0 |
| UniProt | 25,456 | 0 | 0 | 0 | 0 |
| Total | 137,136 | 152,498 | 39,039 | 47,499 | 12,929 |

#### With PharmGKB data

Table 7 summarizes results of the instantiation of PGxO with PharmGKB data (2018-03-05 release).

**Table 7.** Numbers of PGx relationships extracted from PharmGKB v2018-03-05. Some PGx relationships can cause both *Toxicity/ADR* and *Efficacy*.

|  | Caused phenotype |  | Level of evidence |  |  |  |  |  | All |
| --- | --- | --- | --- | --- | --- | --- | --- | --- | --- |
|  | Toxicity/ADR | Efficacy | 1A | 1B | 2A | 2B | 3 | 4 |  |
| # PGx relationships | 1,268 | 1,531 | 44 | 11 | 71 | 97 | 2,270 | 208 | 2,701 |

Figure 3 provides an example of a PharmGKB relationship represented with PGxO. It represents a relationship between the haplotype TPMT*3C and the drug azathioprine. This relationship was extracted with our algorithm named *pharmgkbsql2triples*. Accordingly this algorithm is specified as provenance metadata, along with its version and the version of PharmGKB. This allows coexisting extractions of concurrent versions of PharmGKB or the algorithm. The level of evidence of the relationship is represented with PROV-O concepts.

**Figure 3.**
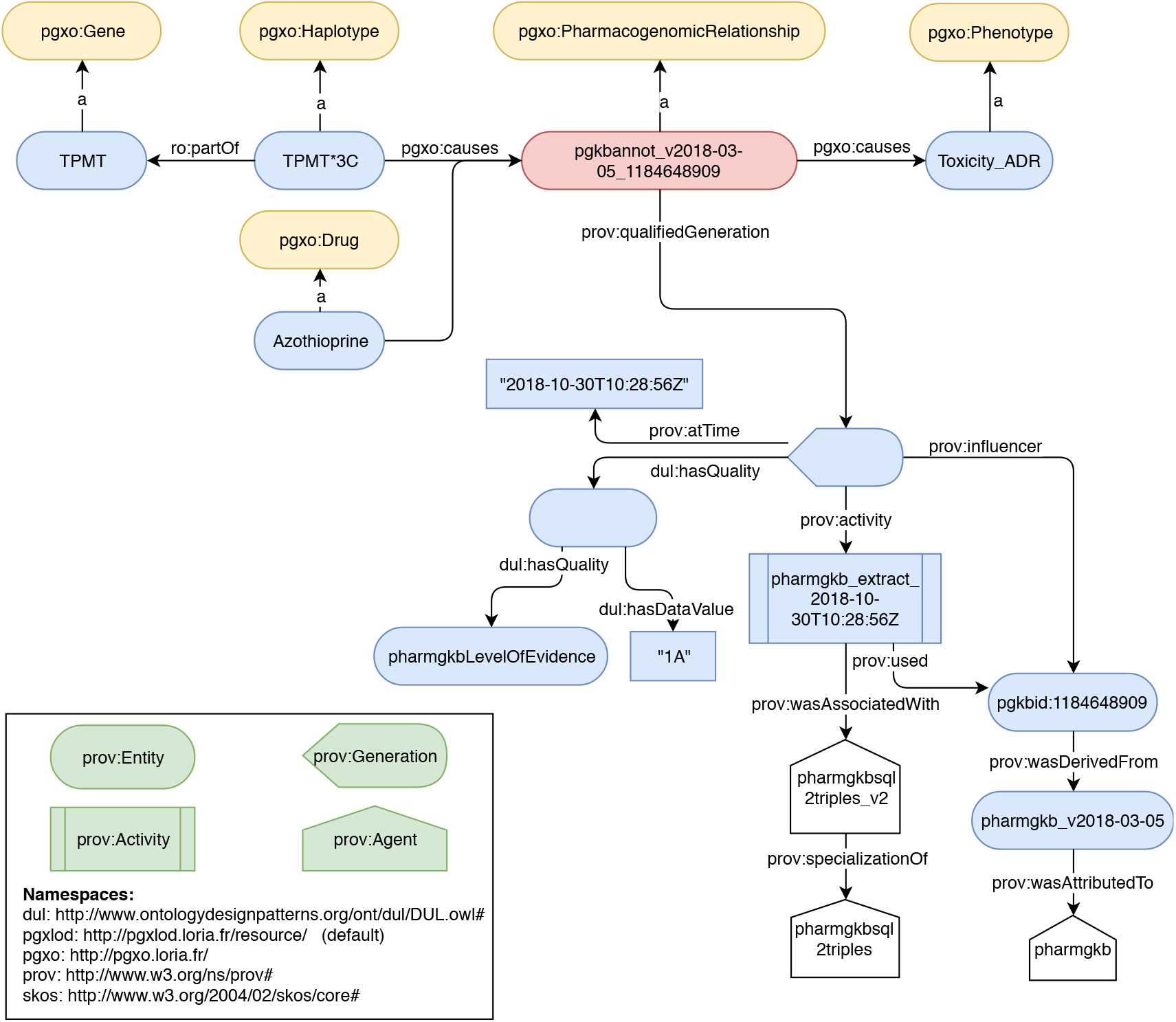
A PGx relationship extracted from PharmGKB on October 30th, 2018 and represented with PGxO. For readability purposes, in some cases labels are used instead of URIs. Only one drug and one variant are represented, whereas this relationship involves more components. The clinical annotation is available at https://www.pharmgkb.org/gene/PA356/clinicalAnnotation/1184648909

#### With the biomedical literature

We instantiated PGxO with the manually annotated sentences of our 307-sentence corpus, and with the output of our prototype for relation extraction on a test corpus formed by the all 176,704 sentences unannotated or annotated. We extracted from these sentences, 51,924 entities (8,412 drugs, 10,812 genes, 8,740 genomic variations and 23,960 phenotypes) and 65,182 PGx relationships between them. Table 8 shows the statistics for the normalization of these entities to identifiers of reference databases or ontologies listed in Table 4. Figure 4 illustrates the RDF encoding of a PGx relationship extracted from the literature. It is noteworthy that the type of relation is encoded in the provenance metadata. Our prototype only outputs relations of the broad type “isAssociatedWith”, but other types could be expected with a more advanced system, e.g. “increases” or “decreases”. Figure 4 also illustrates how the entity representing the TPMT gene reuses an URI from the Bio2RDF transformation of the NCBI Gene database, while the entity representing the 6-mercaptopurine instantiates a MeSH class. This differentiates the use of reference databases or ontologies when normalizing.

**Figure 4.**
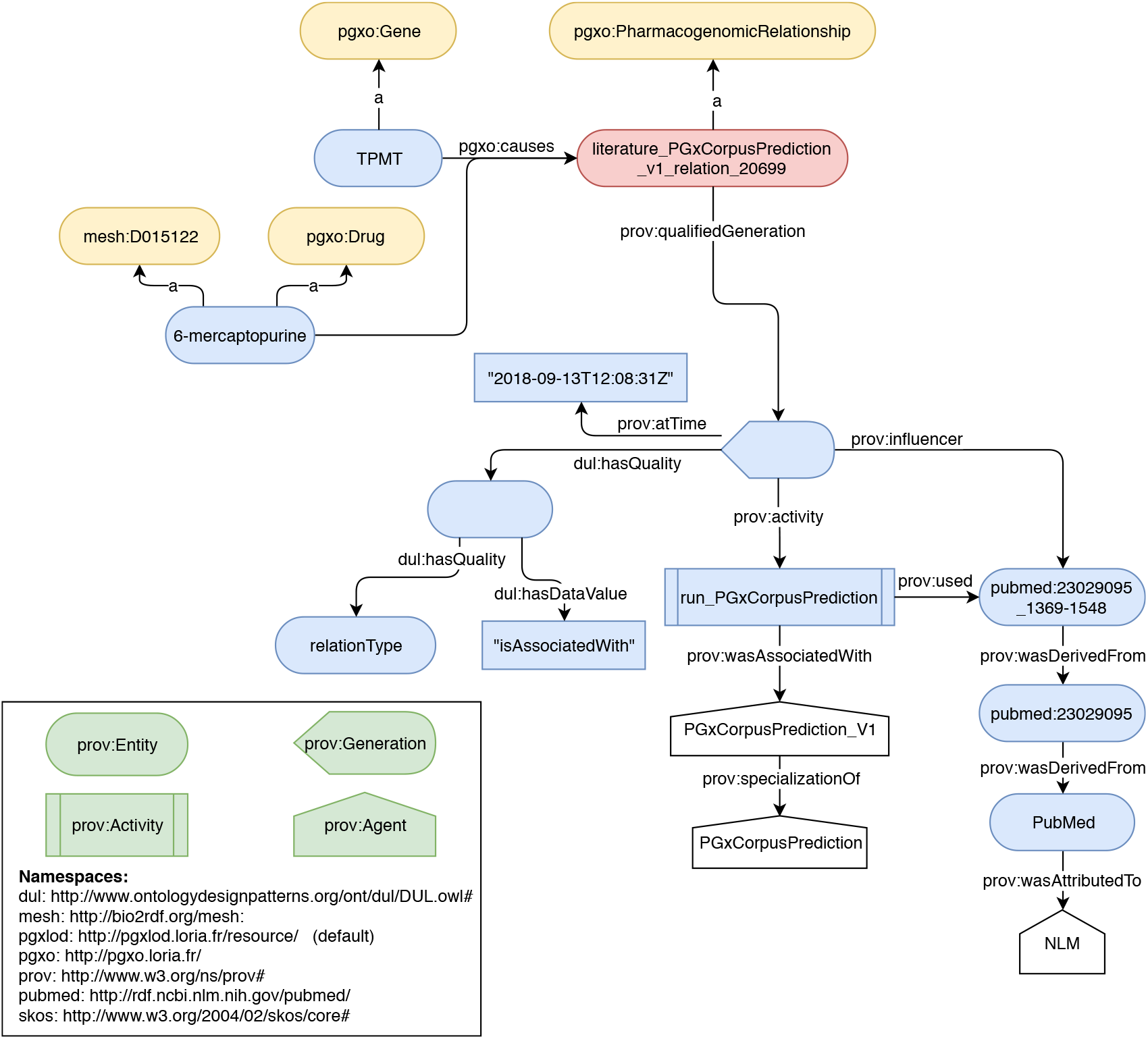
A PGx relationship extracted from the literature on September 13th, 2018 and represented with PGxO. For readability purposes, in some cases labels are used instead of URIs. For example, the TPMT gene is identified with the URI http://bio2rdf.org/ncbigene:7172. The abstract is available at https://www.ncbi.nlm.nih.gov/pubmed/23029095/.

**Table 8.** Numbers of unique entities recognized in the test corpus and successfully mapped with reference databases or ontologies. Reference databases and ontologies are listed in Table 4.

| Database / Ontology | Drug | Gene | GenomicVariation | Phenotype |
| --- | --- | --- | --- | --- |
| MeSH | 1,600 | n/a | n/a | 1,625 |
| ChEBI | 285 | n/a | n/a | n/a |
| ATC | 78 | n/a | n/a | n/a |
| NCBI Gene | n/a | 4,907 | n/a | n/a |
| dbSNP | n/a | n/a | 803 | n/a |
| MEDDRA | n/a | n/a | n/a | 0 |
| PGxLOD | 6,449 | 5,905 | 7,937 | 22,335 |
| Total | 8,412 | 10,812 | 8,740 | 23,960 |

#### With Electronic Health Records and linked biobank studies

Each of the ten studies listed previously results in one instance of a PGx relationship, along with its provenance. Interestingly, out of ten, eight were derived from the BioUV biobank and its linked EHRs [53], one from clinical data of the eMERGE Network [22] and one from data of the HEGP, a

French University Hospital [27]. Out of the same ten relationships, six were obtained from a statistical analysis using linear regression and four using logistic regression. Regarding genetic factors, relationships involve either a single nucleotide polymorphism (7/10), an haplotype (2/10) or an enzyme activity (1/10). For example, Figure 5 represents the instantiation of PGxO, achieved from the results of Neuraz *et al.* [40] and the thiopurine CPIC guidelines. In this particular case, no genetic data was provided in the study, but an enzyme activity. Indeed the TPMT enzyme activity may be considered as a *proxy* for the genotype of the TPMT gene, as stated in the thiopurine-related CPIC guidelines [51]. We added a RDF triple stating that the TPMT activity depends on the TPMT haplotype (with the ro:dependsOn relation type), and documented the provenance of this assertion (with the `pgxo:qualifiedProxy` and `pgxo:qualifiedVariation` relation types and PROV-O concepts and relation types). This representation is possible because our axioms defining a PGx relationship allow using a phenotype as a proxy for a genetic factor instead of a genetic factor itself. We expect this to facilitate later comparison of the results of studies without genetic data, to the state of the art.

**Figure 5.**
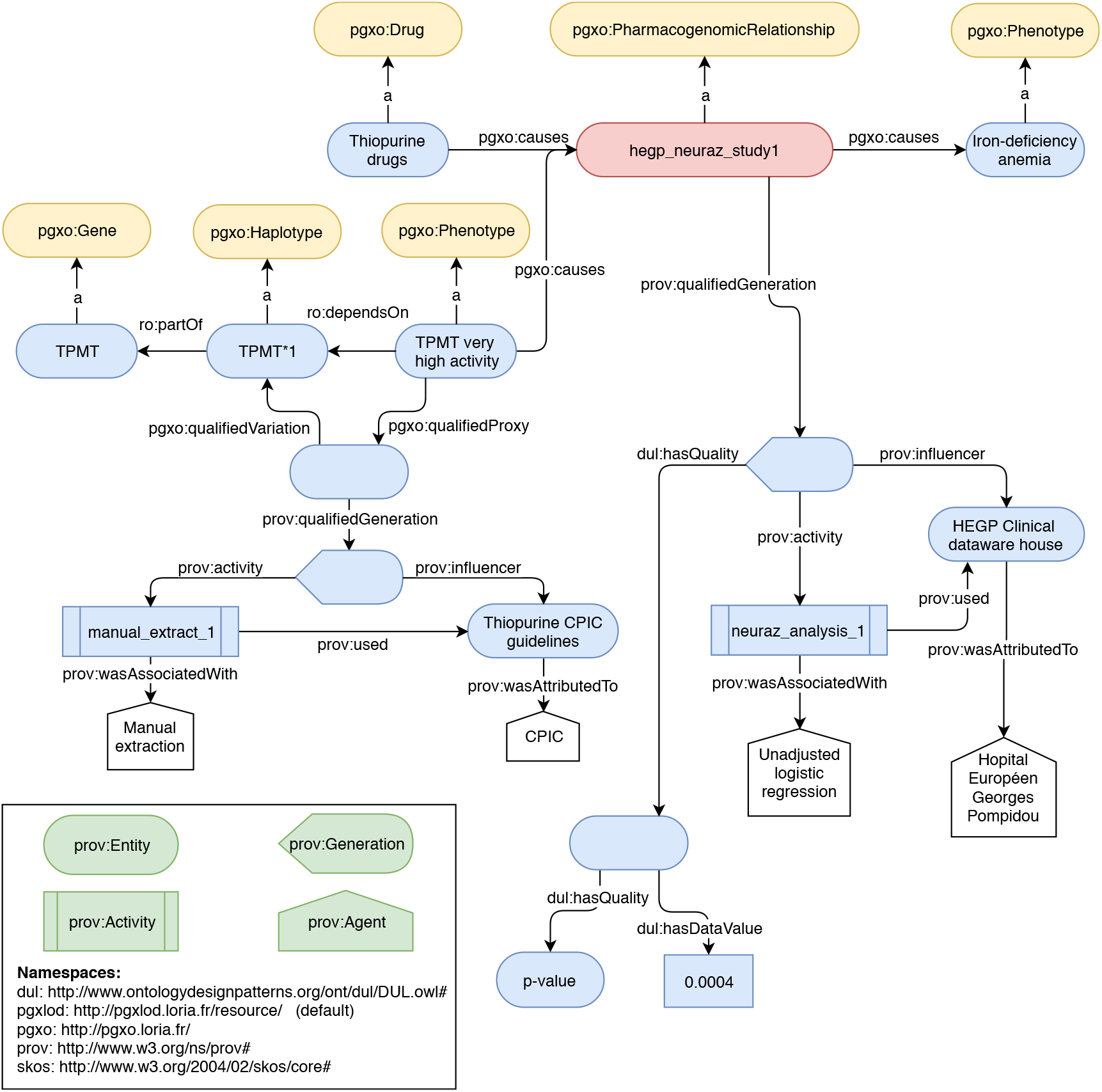
A PGx relationship discovered from EHRs [40] and represented with PGxO. The initial association discovered from EHRs is standing between a drug response and the TPMT activity, i.e. a phenotype. The later is considered a *proxy* to the genotype of the TPMT gene, as stated by the CPIC guidelines. For readability purposes, in some cases labels are used instead of URIs.

### Reconciliation rules

#### Definition and implementation of the reconciliation rules

We defined five rules for simple pair-wise comparison of PGx relationships. These rules are able to identify when two PGx relationships with distinct provenances are in fact referring to the same knowledge unit, to a more specific knowledge unit, or to related knowledge units (to some extent). Indeed, among the five rules, Rule 1 is dedicated to identifying identical relationships, Rules 2, 3, and 4 to identifying broader/narrower ones and Rule 5 to identifying PGx relationships related by some of their components. All five rules are provided in Additional file 1 as well as examples of their application on RDF graphs. In the next paragraphs, as an example, the simpler rule, Rule 1, identifying when two PGx relationships are referring to the same knowledge unit is presented and illustrated. Other rules are a bit more complex, but follow the same principles.

Rules compare PGx relationships on the basis of their components, i.e. sets of drugs, genetic factors and phenotypes. Accordingly, considering r, an instance of the PharmacogenomicRelationship concept from a knowledge base *𝒦𝓑*, the following sets are defined.

**Notation 1.** *We denote D, the set of instances of* Drug *that cause* **r***, defined as:*

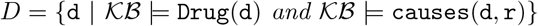

**Notation 2.** *We denote G, the set of instances of* GeneticFactor *that cause* **r***, defined as:*

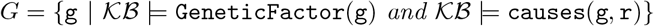

**Notation 3.** *We denote P, the set of instances of* Phenotype *caused by* **r***, defined as:*

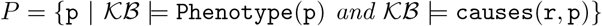

Therefore, when comparing two PGx relationships denoted by r_1_ and r_2_, the sets of their components will be denoted *D*_1_, *G*_1_, *P*_1_ and *D*_2_, *G*_2_, *P*_2_. The first reconciliation rule identifies when two PGx relationships are referring to the same knowledge unit; it is defined as follows:

**Rule 1.** *D*_1_ = *D*_2_ AND *G*_1_ = *G*_2_ AND *P*_1_ = *P*_2_ ⇒ owl:sameAs(r_1_, r_2_)

This rule states that when two relationships involve the same sets of drugs, of genetic factors and of phenotypes, they refer to the same knowledge unit. Therefore, the link owl:sameAs(r_1_, r_2_) should be added to the knowledge base. For example, consider the RDF graph presented in Figure 6. We have:

- *D*_1_ = *D*_2_ = `{warfarin}`
- *G*_1_ = *G*_2_ = `{CYP2C9}`
- *P*_1_ = *P*_2_ = `{cardiovascular diseases inst1}`

**Figure 6.**
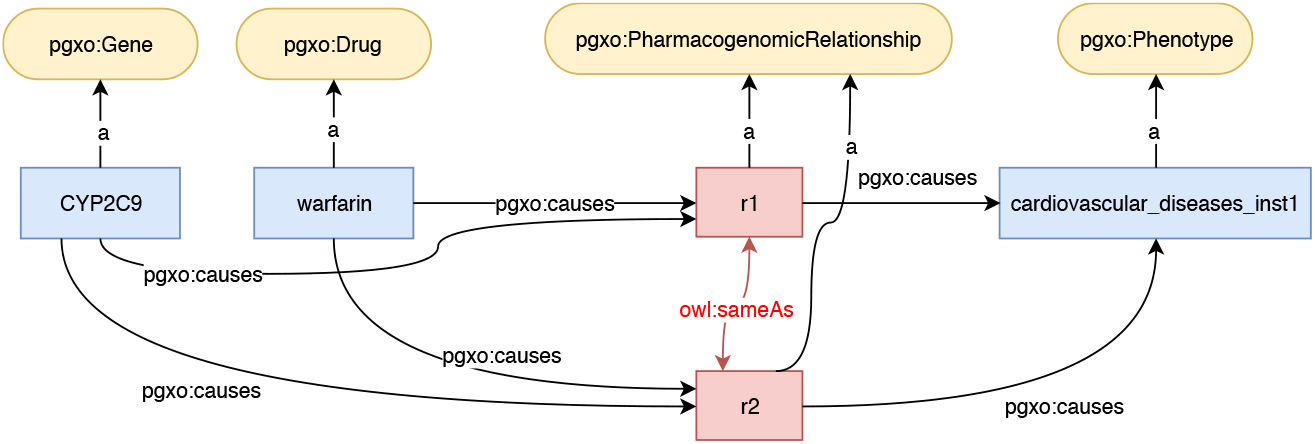
Example of a RDF graph on which a reconciliation rule identifies that two PGx relationships are identical. The owl:sameAs links result of the application of the rule.

Therefore, the left part of Rule 1 is true, and the link owl:sameAs(r_1_, r_2_) should be added to the knowledge base.

Other rules, details and examples are available in Additional file 1. Rules 2, 3 and 4 conclude in indicating that a relationship is more specific than the other by adding the link skos:broadMatch(r_1_, r_2_) to the knowledge base. Rule 5 concludes that they are related by adding the link skos:relatedMatch(r_1_, r_2_).

#### Execution of the reconciliation rules on PGxLOD

We executed our reconciliation rules on PGxLOD v2 (Table 5). As each relationship is compared to all the others, this led to 68, 430 × 68, 431 = 4, 682, 733, 330 comparisons performed.

This execution generated owl:sameAs links (Table 9), `skos:broadMatch` links (Table 10) and `skos: relatedMatch` links between the PGx relationships in PGxLOD. We observed skos:relatedMatch (37,758) only between pairs of relationships extracted from the literature. Interestingly, for 132 PGx relationships from PharmGKB an identical relationship was found (generating `132 owl:sameAs` links). Additionally, 14 sentences from the biomedical literature are identified as more generic than what is stated in the EHR+biobank studies.

**Table 9.** Number of `owl:sameAs` links between PGx relationships from each source.

|  | EHRs | Literature | PharmGKB |
| --- | --- | --- | --- |
| EHRs | 0 | 0 | 0 |
| Literature | 0 | 109,078 | 0 |
| PharmGKB | 0 | 0 | 132 |

**Table 10.**
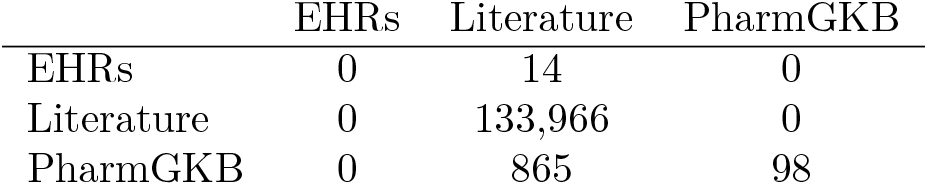
Number of `skos:broadMatch` links between PGx relationships from each source. Rows represent origins of the links and columns represent destinations.

|  | EHRs | Literature | PharmGKB |
| --- | --- | --- | --- |
| EHRs | 0 | 14 | 0 |
| Literature | 0 | 133,966 | 0 |
| PharmGKB | 0 | 865 | 98 |

We can notice that skos:broadMatch links exist between different sources while `owl:sameAs and skos:relatedMatch` links only refer to PGx relationships from the same sources. Some possible explanations reside in a lack of mappings between entities in different vocabularies and the use of broad phenotypes for PGx relationships from PharmGKB (i.e. *Toxicity/ADR*, *Efficacy*) that are distinct from more specific phenotypes elsewhere stated such as, for example, *cardiovascular diseases*.

## Discussion

Instantiating PGxO with knowledge extracted from various sources allows to answer the defined *competency questions* : we are able to represent PGx relationships extracted either from the state of the art (reference databases or the literature) as well as from EHR+biobank studies. The use of heterogeneous sources for instantiating our ontology improved in several ways the modeling of PGx relationships previously drafted in [37]. Among other things, we enabled the representation of phenotypes as proxies for a specific genotype, such as an enzyme activity. The encoding of metadata has also been enriched to enable the encoding of the various metrics associated with the different kinds of knowledge extractions.

By using Semantic Web technologies, our global framework for knowledge comparison in PGx can easily leverage knowledge defined elsewhere such as ontologies or other available LOD sets. This is of particular importance as the reconciliation rules depend on existing mappings and subsumption relations. Moreover, the proposed encoding can easily evolve depending on one’s needs. However, in a data warehousing perspective, Semantic Web technologies require high data maintenance to follow the evolution of associated databases, LOD sets and ontologies. Therefore, one challenge is to keep PGxLOD up-to-date with respect to the associated data sources.

Several directions are considered to continue this work. Regarding the extraction from PharmGKB, more detailed drug response phenotypes could be extracted from the plain-text sentences describing the clinical annotations in the database. This would require text mining similar to what we have done for processing literature but this would enable a more accurate comparison between the content of PharmGKB and other sources.

Our prototype for knowledge extraction from the literature constitutes solely a baseline. It faces limitations relatively easy to improve. First, the NER model, in its current form, does not detect discontinuous entities that may appear in the literature (such as “the *response of* the selective serotonin reuptake inhibitors *paroxetine*” where the entity “response of paroxetine” is discontinuous). This is a limitation since missed entities lead to missed relationships. In addition, the two steps NER procedure can only detect fairly simple included entities. In practice, multiple levels of inclusion can be observed in the literature and cannot be captured by our system. Finally, a larger training corpus would improve the performance of the learned models, since deep learning architectures usually require large annotated corpora in order to achieve reasonable performances. In addition, the normalization process, which results are reported in Table 8, is naive and may be improved using lexical resources and ontology repositories. Those list term synonyms, variants and bridges between these resources.

The manual instantiation of PGxO with knowledge extracted from EHRs constitutes only a proof of concept. One notable drawback is that gene variants and precise drug response phenotypes are not available in most cases. Thus, the knowledge discovery process needs to rely on proxies such as a phenotype being a marker of the patient genotype or a stable dose requirement being a marker of the patient sensitivity to the considered drug. Therefore, a PGx relationship discovery from EHRs would benefit from a more complete list of proxies. To our knowledge, no such list is available. In addition, more contextual information about knowledge discovered from patient data would be of interest. For example, the ethnicity of patients [29] or the indications for which patients are treated [14] may be necessary to properly document some PGx relationships. Considering these challenges, one perspective of the current work relies in automatically instantiating PGxO with knowledge extracted by mining EHRs.

The proposed reconciliation rules were executed on PGxLOD, providing first results of reconciliation. However, to compare PGx relationships involving entities from different vocabularies, the rules rely on the existence of equivalence or subsumption relationships between the URIs of these entities. Therefore, a major task and perspective resides in completing the mappings between entities of various provenances. Using both concept hierarchies and ontology-to-ontology mappings defined in the UMLS [6] or the NCBO Bioportal [25] may improve knowledge comparison. Especially, this may be particularly useful when considering knowledge extracted from EHRs, which are expressed with concepts of ontologies used in the encoding of clinical practice such as ICD or RxNorm. Finally, the reconciliation rules strictly compare the components of a PGx relationships: drugs, genetic factors, and phenotypes. However, other features could be considered, such as the specific chemicals of a drug. Such features could be involved in a fuzzy comparison highlighting similar but not strictly equivalent relationships.

## Conclusions

In this article, we presented a simple ontology called PGxO to represent pharmacogenomic knowledge and its provenance. With the combined use of PROV-O and DUL, we demonstrated that PGxO can structure knowledge extracted from various sources such as reference databases (i.e. PharmGKB), the literature, clinical guidelines or EHR+biobank studies. We also defined and implemented a set of rules allowing to compare and reconcile PGx knowledge units from different sources. PGxO and the reconciliation rules constitute a first step to a semantic framework able to represent, trace, confront and reconcile PGx relationships from various origins. A first experiment with these rules highlights equivalent and comparable pieces of knowledge across various data sources, opening perspectives for fine grained comparison and interpretation of the content of PGx sources. Finally, we think that the resulting and integrated data set called PGxLOD constitutes by itself a valuable resource for PGx research. This data set is made available to the community and will be improved with additional knowledge from the state of the art and from EHR mining.

## Additional Files

### Additional file 1 — PGxO reconciliation rules

This PDF file provides the definition of the five reconciliation rules and illustrates their behavior with concrete examples.

## Author’s contributions

PM, AN and AC designed the experiments. PM, JL and AC wrote the manuscript. PM, CJ and AC designed the ontology and the encoding of the metadata. PM, AN and AC designed the reconciliation rules. PM implemented and ran the process of knowledge reconciliation. PM and GH implemented and ran the processes of knowledge extraction from PharmGKB. JL implemented and ran the processes of knowledge extraction from the biomedical literature. AC extracted knowledge related to biobanks and EHRs studies. PR set up and administers the virtuoso server for PGxLOD. AT and CJ advised on the definition of mappings between LOD entities. All authors read and approved the final manuscript.

## Funding

This work is supported by the *PractiKPharma* project, founded by the French National Research Agency (ANR) under Grant No. ANR-15-CE23-0028, by the IDEX “Lorraine Université d’Excellence” (15-IDEX-0004), by the *Snowball* Inria Associate Team, and by the European Union’s Horizon 2020 research and innovation program under the Marie Sklodowska-Curie grant agreement No. 701771.

## Acknowledgments

We acknowledge the participants of the NETTAB 2017 workshop for their constructive feedback on the preliminary results of this work. This work was conducted using the Protégé resource, which is supported by grant GM10331601 from the National Institute of General Medical Sciences of the United States National Institutes of Health.

